# Impact of pathogen genetics on clinical phenotypes in a population of *Talaromyces marneffei* from Vietnam

**DOI:** 10.1101/2023.03.30.534926

**Authors:** Poppy Sephton-Clark, Thu Nguyen, Ngo Thi Hoa, Philip Ashton, H. Rogier van Doorn, Vo Trieu Ly, Thuy Le, Christina A. Cuomo

## Abstract

Talaromycosis, a severe and invasive fungal infection caused by *Talaromyces marneffei*, is difficult to treat and impacts those living in endemic regions of southeast Asia, India, and China. While 30% of infections result in mortality, our understanding of the genetic basis of pathogenesis for this fungus is limited. To address this, we apply population genomics and genome wide association study approaches to a cohort of 336 *T. marneffei* isolates collected from patients who enrolled in the Itraconazole versus Amphotericin B for Talaromycosis (IVAP) trial in Vietnam. We find that isolates from northern and southern Vietnam form two distinct geographical clades, with isolates from southern Vietnam associated with increased disease severity. Leveraging longitudinal isolates, we identify multiple instances of disease relapse linked to unrelated strains, highlighting the potential for multi-strain infections. In more frequent cases of persistent talaromycosis caused by the same strain, we identify variants arising over the course of patient infections that impact genes predicted to function in the regulation of gene expression and secondary metabolite production. By combining genetic variant data with patient metadata for all 336 isolates, we identify pathogen variants significantly associated with multiple clinical phenotypes. In addition, we identify genes and genomic regions under selection across both clades, highlighting loci undergoing rapid evolution, potentially in response to external pressures. With this combination of approaches, we identify links between pathogen genetics and patient outcomes and identify genomic regions that are altered during *T. marneffei* infection, providing an initial view of how pathogen genetics affects disease outcomes.

## Introduction

The thermally dimorphic fungal pathogen, *Talaromyces marneffei*, causes talaromycosis, a severe fungal infection that effects immunocompromised people living in, or traveling to, endemic regions spanning Southeast Asia, India, and China (1, 2). People with advanced HIV disease are particularly affected, in whom the prevalence of talaromycosis can reach 26.5% in some regions (1). The impact of this disease is an estimated 4,900 (95% CI 2500–7300) deaths annually (2). Talaromycosis is difficult to treat, with mortality rates of up to 30% (3, 4), and rates of infection increase throughout the rainy season, disproportionately impacting farmers and agricultural workers (2, 5).

*T. marneffei* infection originates primarily through inhalation (6). Once in the lungs, *T. marneffei* can disseminate to multiple organs, including the liver, spleen, lymph nodes, blood, bone marrow, bone and skin (6–9). The infectious propagule includes aerosolized conidia, which may transition to the pathogenic yeast form upon inhalation (10). Effective containment by lung-resident primary immune cells is critical to preventing dissemination, as *T. marneffei* possesses multiple virulence traits that enable its persistence within the human host. These virulence factors include the ability to grow at 37 ℃ (9), the production of reactive oxygen species (ROS) detoxifying enzymes that may promote survival within host macrophages (11), the secreted galactomannoprotein Mp1 that suppresses the proinflammatory host immune responses (12), and multiple laccases important for resisting external stressors and killing by phagocytes (13). Further identification of genes important for virulence in *T. marneffei* is still needed to enable a better understanding of how *Talaromyces* adapts and survives within the host.

Recently generated complete genome assemblies for representative isolates of *T. marneffei* from north and south Vietnam (14) will enable further discovery of genes essential for virulence, as well as the evaluation of genetic variability across clinical isolates. Genetic variation in populations of many pathogenic fungi has been surveyed (15–19), and our initial understanding of variation across populations of *T. marneffei* has been guided by work that uncovered the population delineation of country-linked clades, based on a multilocus sequence typing (MLST) study (20). This prior work supplied an overview of the population structure of *T. marneffei* and provided a foundation for further analysis, based on whole genome sequencing, to better understanding the genetic variation within clinical populations of *T. marneffei*. Characterizing genetic heterogeneity in populations of *Aspergillus*, *Cryptococcus*, and *Candida* has enabled a better understanding of resistance mechanisms and virulence traits through the study of aneuploidy (21–23), copy number variation (17), and variant-phenotype association studies (15, 19, 24, 25). Population studies leveraging whole genome data have also allowed for the discovery of clade specific adaptation and evolution, such as the loss of subtelomeric adhesins in the only non-outbreak causing *Candida auris* clade (clade II) (26). In addition to understanding heterogeneity across populations, these approaches have enabled the interrogation of variability evolving in isolates over time, through the study of patient longitudinal and relapse isolates (27–29), and microevolution of isolates passaged through infection models (30–32).

In this study, we leverage a cohort of 336 clinical *T. marneffei* isolates from northern and southern Vietnam (33) to explore population diversity, determine levels of recombination, identify genomic regions under selection, and assess genomic predictors of clinical phenotypes. With this approach, we identify two geographically distinct populations showing similar levels of recombination within and between clades. Notably, we find that clinical aspects of disease severity differ between isolates from these two clades. We employ longitudinal samples isolated from blood and skin lesions to identify variants arising over the course of invasive infection. We observe multiple instances of disease recrudescence/relapse that are linked to the independent introduction of an unrelated isolate, in addition to the more frequent detection of persistent infection by the same isolate. We identify multiple genomic regions that appear to be under selection across these two populations, with copy number variation arising throughout the population, and multiple genetic variants showing a significant association with relevant clinical parameters including a high initial fungal burden in the blood, mortality, relapse, immune reconstitution inflammatory syndrome (IRIS), and poor treatment response.

## Materials and Methods

### Sample collection and antifungal susceptibility testing

The 336 clinical *T. marneffei* isolates were obtained from cultures of blood and/or skin lesions of 272 patients from 5 hospitals across Vietnam who participated in the Itraconazole versus Amphotericin B for Talaromycosis (IVAP) trial conducted between October 2012 and December 2016 in Vietnam (33). The *in vitro* antifungal susceptibility of all 336 *T. marneffei* yeast isolates against itraconazole and amphotericin B was determined using the M27-A3 reference method for testing yeasts as per the clinical and laboratory standards institute (CLSI) (34). The minimum inhibitory concentration (MIC) was determined by both visual and optical endpoints with growth inhibition of 90% for amphotericin B and 50% for itraconazole according to the CLSI. The study was approved by the ethics and scientific committees of the University of Oxford, all five participating hospitals in Vietnam, and Duke University. All patients gave informed consent for their *T. marneffei* clinical isolates to be stored and used for this research.

### Clinical metadata

De-identified clinical metadata was collected and shared by the trial investigators. These data included presence of fever, measures of baseline fungal burden in colony forming units (CFUs) per ml of blood, rates of fungal clearance in CFUs/ml/day, 24-week mortality, time to clinical resolution (defined as resolution of fever, skin lesions, and positive blood culture), incidence of relapse and immune reconstitution inflammatory syndrome (IRIS), and respiratory failure over 24 weeks of follow-up. These patient outcome variables were originally defined in the IVAP trial (33). All statistical analyses were performed in R 3.6.0.

### Genomic DNA Isolation and sequencing

DNA was purified using the yeast genomic DNA purification kit (VWR). Briefly, cells were disrupted with sodium dodecyl sulphate (SDS) (VWR) at 65 °C. The pellet was then treated with ammonium acetate and RNase A (VWR). DNA was precipitated with isopropanol and stored in Tris-EDTA buffer at a pH of 8.0 (VWR). DNA purity was assessed via Nanodrop (Thermo Fisher Scientific) and 1% agarose gel electrophoresis. The DNA concentration was measured via Qubit fluorometer (Thermo Fisher Scientific). Initial library construction for the isolates entailed tagmentation via the Nextera XT DNA library preparation protocol (Illumina). Isolates were initially sequenced on a HiSeq 3000 (Illumina), generating 100bp paired reads. Samples with poor coverage resulting from this first method then underwent additional sequencing, with libraries generated in accordance with the NEBNext Ultra II DNA Library Prep protocol (New England Biolabs). Libraries were sequenced on a HiSeq X10 (Illumina), generating 150bp paired reads (minimum average coverage 10X).

### Variant analysis

To identify variants for all samples, reads were aligned to the *T. marneffei* (11CN-03-130) reference genome (GCA_009650675.1) (14) with BWA-MEM version 0.7.17 (35). GATK v4 variant calling was carried out as documented in our publicly available cloud-based pipeline (36) (https://github.com/broadinstitute/fungal-wdl/tree/master/gatk4). Post calling, variants were filtered on the following parameters: QD < 2.0, QUAL < 30.0, SOR > 3.0, FS > 60.0 (indels > 200), MQ < 40.0, GQ < 50, alternate allele percentage = 0.8, DP < 10. Variants were annotated with SNPeff, version 4.3t (37).

### Population genomics and genome wide association studies

A maximum likelihood phylogeny was estimated using 281,773 segregating SNP sites present in one or more isolates, allowing ambiguity in a maximum of 10% of samples, with RAxML version 8.2.12 GTRCAT rapid bootstrapping (38). The phylogeny generated was visualized with ggtree (39). These SNP sites were also used to compute an unrooted phylogenetic network with splitstree4 (40). To assess recombination, linkage disequilibrium (LD) decay was estimated with vcftools version 0.1.16, calculating LD for 1000bp windows, with a minimum minor allele frequency of 0.1, and the --hap-r2 option (41). To identify regions of the genome under selection within these populations, PopGenome 2.7.5 was used to perform composite likelihood ratio analysis, assess nucleotide diversity and divergence, and calculate Tajima’s D and F-Statistic scores (per chromosome, by 5kb windows) (42). Copy number variation was calculated with CNVnator v0.3 (43). Association analysis between clinical data, *in vitro* phenotypes, and variants was carried out using PLINK v1.08p formatted files and Gemma version 0.94.1 (44) (options: centered relatedness matrix gk 1, linear mixed model), as previously described (15).

### Centromere identification

To identify centromere locations, the GC% was calculated across individual chromosomes with a sliding window approach in R 3.6.0. Repeat motifs across the proposed centromere regions were identified with RepeatModeler 2.0 (45). Recombination rates across individual chromosomes were estimated with a random subset of 50 isolates spanning north and southern clades, via Ldhelmet 1.10 per the best practices workflow (46).

## Results

### Characterization of genome structure

For this study, we leveraged a complete *T. marneffei* genome generated from isolate 11-CN-03-130. This assembly consists of 8 chromosomes, with telomeric sequences present at both ends of all chromosomes excepting one ending in ribosomal DNA repeats (14). This haploid genome encodes 10,025 genes, including rRNA content on chromosome 3, with a BUSCO completeness score of 97.7% (14). To determine the location of the centromeric regions of these chromosomes, we calculated GC% across all chromosomes and detected single regions displaying notable reductions in GC% for each chromosome, corresponding to likely centromeric positions (**Figure 1a**). To determine the composition of these proposed centromeric regions, we identified repetitive sequences based both on similarity to known elements and de novo self-alignments, identifying long interspersed nuclear elements (LINE) and DNA/TcMar-Fot1 transposons throughout these putative centromeric regions (**Figure 1b**). The repetitive content across these regions is consistent with the repetitive nature of centromere regions observed across other species of filamentous fungi (47). These proposed regions range from 15.5-28 kb in length, with an average centromere length for all chromosomes of 23 kb. To assess whether rates of recombination across these proposed centromeric regions are lower when compared to the non-centromeric regions, we calculated recombination rates per chromosome, with variant information from a subset of 50 isolates sequenced in this study. We saw similar rates of recombination across chromosomes (**Figure S1**), and while we did not observe any characteristic dips in recombination rate that might be associated with centromeric locations, the proposed centromeric regions displayed rates that were in line with those observed across the entire chromosome.

**Figure 1.**
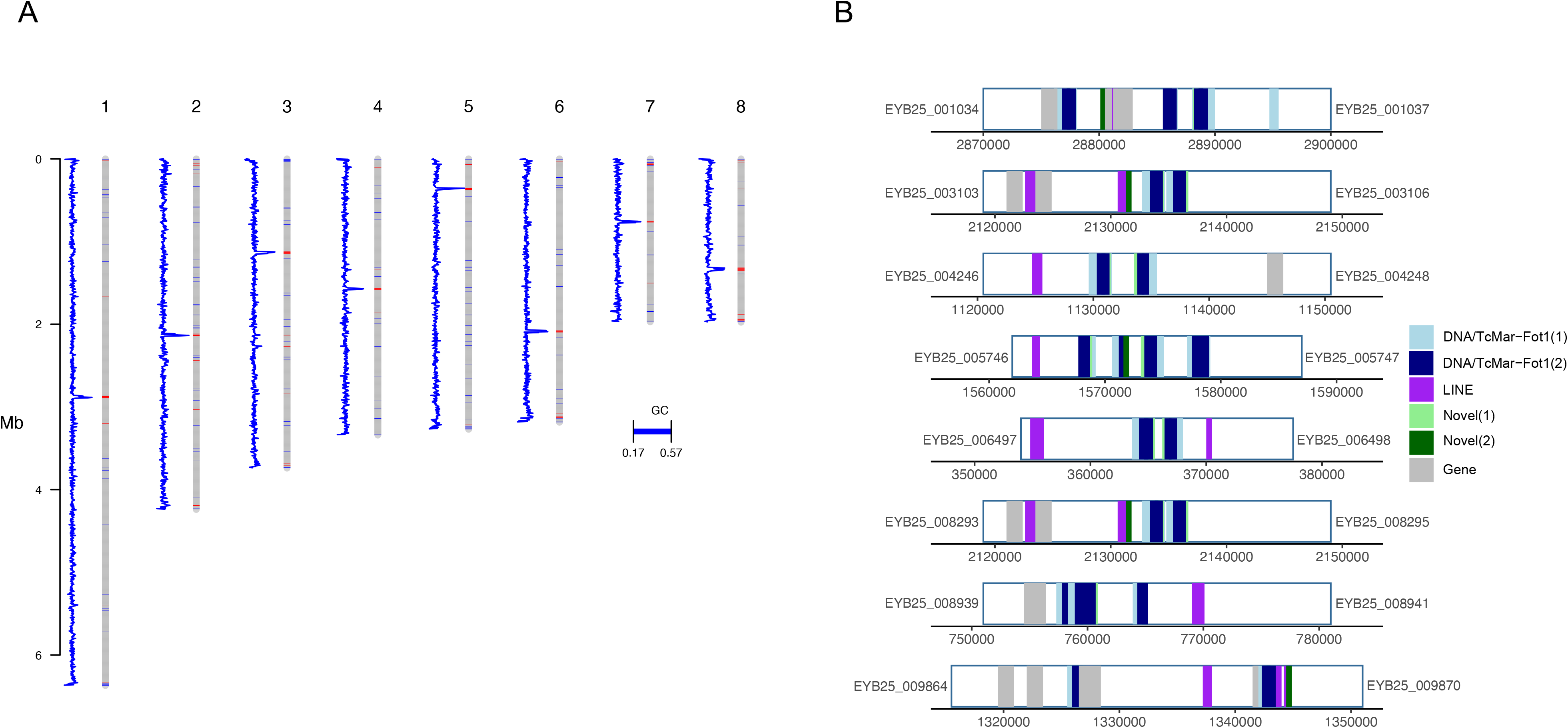
*T. marneffei* centromere identification. A sliding window analysis of GC% and repetitive element content was used to predict candidate centromere locations. A) GC% for each chromosome (blue line plot), calculated and plotted in 500bp windows beside each chromosome (grey bars). Regions of low GC (17-40%) are shown in red on each chromosome. B) Proposed centromere regions with repetitive elements colored by class (highlighted in blue, purple, and green), protein-coding genes shown in grey, with the flanking gene IDs listed either side (EYB25 prefix).

### Population structure and geographical clades

To finely characterize genetic variation of clinical populations, we compared the genomes of *T. marneffei* isolated from HIV-positive patients enrolled in the IVAP trial across northern and southern Vietnam (33). For all individuals, *T. marneffei* isolates were obtained from blood cultures taken at enrollment (day 0). For a subset of individuals, *T. marneffei* isolates were obtained from both cultures taken at enrollment (day 0), and from longitudinal time points for both blood and skin sites. Longitudinal samples were collected between day 4 and day 240, with the median longitudinal sample collection time being 1 month. We performed whole genome sequencing on all isolates collected, requiring a minimum genome coverage of 10X. We then called variants for these isolates, using a reference genome assembly (14) of isolate 11CN-03-130, also collected from southern Vietnam as part of the IVAP trial.

To assess population structure, we generated a maximum likelihood phylogeny from segregating SNP site information (**Figure 2**). Isolates clearly separate into two clades representing the northern and southern regions of Vietnam from which they were isolated. This pattern is consistent with prior geographical substructure with clade separation based on region of collection (20). Clinical and *in vitro* susceptibility phenotypes appear well distributed across the phylogeny for these isolates (**Figure 2, Figure S2**). Northern isolates show slightly higher levels of diversity (**Table 1**), harboring alleles with a higher number of SNPs, and displaying significantly longer terminal branch lengths than southern clade isolates.

**Figure 2.**
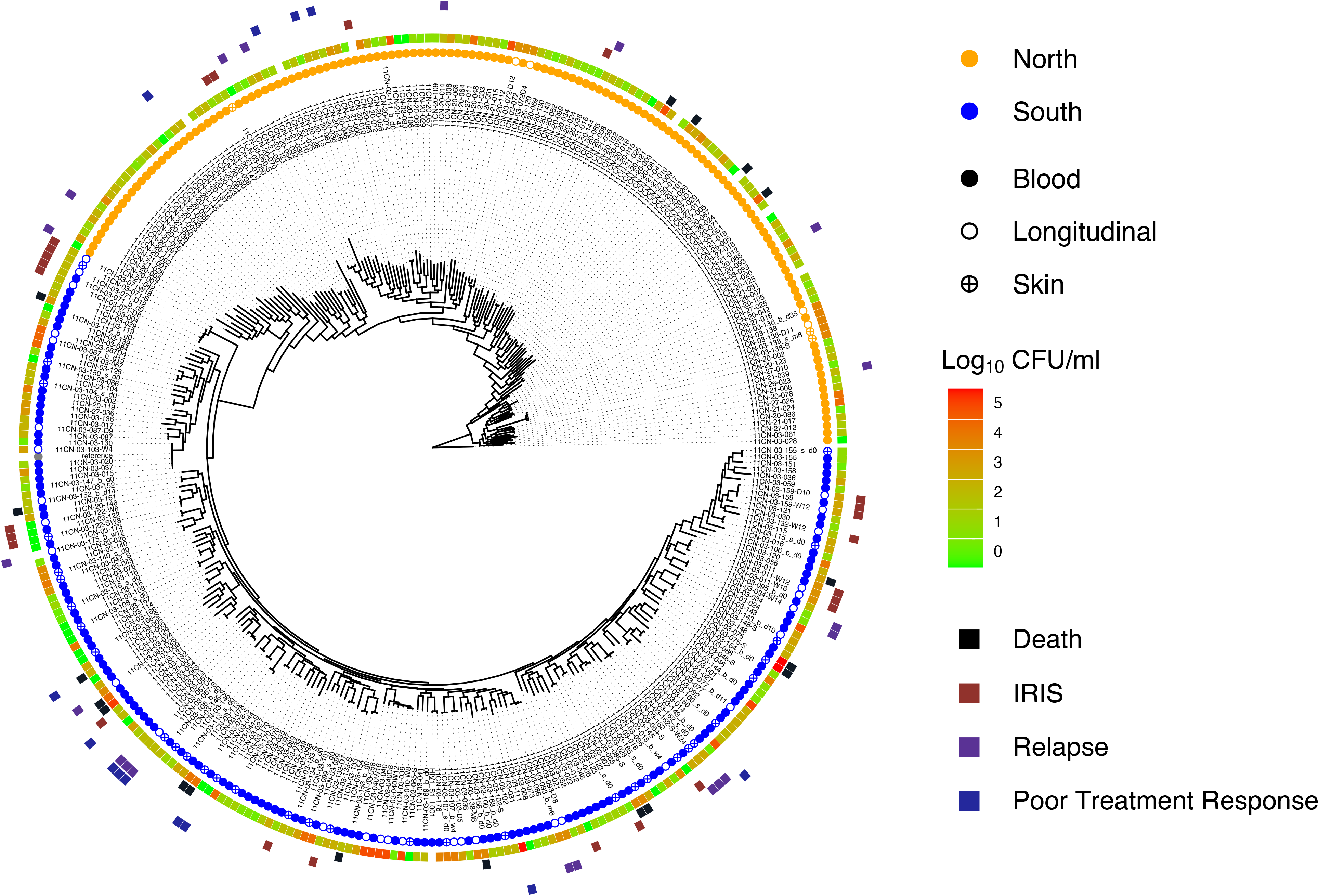
Phylogeny of patient isolates reveals no genetic clusters of severe outcomes. A maximum likelihood phylogeny was estimated using segregating SNP sites. Isolates separate distinctly into northern and southern clades, with both clades having 100% bootstrap support. The colored circles correspond to clade and sample type. Metadata for the clinical isolates includes initial fungal burden for a patient, indicated by the Log_10_ CFU/ml heatmap, and patient outcomes that are indicated by colored squares in the outer perimeter of the phylogeny.

**Table 1.** Genomic and population characteristics of isolates sampled.

|  | northern Vietnam | southern Vietnam |
| --- | --- | --- |
| Sample size (baseline blood, skin, and longitudinal) | 141 | 195 |
| LD50 | 12kb | 12kb |
| $\pi$ | 0.00077 | 0.00051 |

To assess recombination levels between and across these two clades, we calculated linkage disequilibrium (LD) decay for the northern clade, the southern clade, and all isolates combined (**Figure S3, Table 1**). We see similar rates of decay, and identical LD50 values (12kb) across all groups, indicating that neither clade is recombining exclusively within their group.

### Clinical outcomes and longitudinal samples

To assess whether clinical phenotypes or *in vitro* antifungal susceptibility values significantly differed by clade, we performed a series of statistical analyses to assess associations between clade and phenotype. We did not observe statistically significant differences between clades in considering 24-week mortality, rates of relapse, IRIS, or MICs of itraconazole and amphotericin B. However, some striking differences were detected. While isolates from the southern clade have comparable rates of fungal clearance to isolates from the northern clade (**Figure S4a**), the southern isolates exhibit significantly higher baseline fungal burden (**Figure S4b**) (Wilcoxon rank sum test, p=0.024), significantly higher prevalence of fever (Fisher exact probability test, p=0.0002), and significantly decreased incidence rates of respiratory failure (Fisher exact probability test, 0.0021) (**Table 2**). Given the increased presence of fever and baseline fungal burden across isolates from the southern clade, we sought to assess whether additional factors including patient age, antiretroviral (ARV) treatment status or CD4 counts were confounders in these analyses. For patients included in this analysis, the median age was 33, the median CD4 count was 10, and 44% of patients had previously received ARV treatment. Neither age, ARV treatment status, or CD4 counts were significantly correlated with the presence of fever or baseline fungal burden, indicating that isolate clade is a contributing factor to some aspects of disease severity in patients from Vietnam.

**Table 2.** Frequency of clinical phenotypes associated with northern and southern clades.

| Clinical phenotypes | northern Vietnam | southern Vietnam | P value<sup>a</sup> |
| --- | --- | --- | --- |
| Median Baseline fungal burden (Log<sub>10</sub> CFU/ml) | 2.1 | 2.5 | 0.024 |
| Fever | 10 | 34 | 0.0002 |
| Respiratory failure | 45 | 24 | 0.0021 |

<sup>a</sup>Fishers exact test and Wilcoxon rank sum test were used to calculate P values.

To assess the genetic relationships between isolates sampled from different body sites (skin vs blood) of a patient at the same time point, and isolates sampled at different time points from the same patient (pre vs post-treatment), we calculated the number of SNP differences between each isolate. Over 80% of isolates sampled at longitudinal time points show a small number of SNP differences (≤10) when compared to their baseline counterparts (**Table 3**). Some of the genes that were impacted by SNPs arising during infection include a predicted Myb-like binding domain (EYB25_007616), a transcription factor (EYB25_005954), and AMP-binding protein (EYB25_000740), an ORC DNA replication protein (EYB25_004486), and a cytochrome P450 (EYB25_003095). We performed additional tests to assess differences in the MICs between incident and post-drug exposure isolates. We observed a general trend of longitudinal isolates maintaining (62%) or increasing (12%) MIC values over time when compared to their baseline counterparts.

**Table 3.** Genomic changes between related isolates taken from the same patients over time, or from different body sites at the same time point.

| Baseline | Longitudinal | Time (days) | SNPs | Gene impacted |
| --- | --- | --- | --- | --- |
| 11CN-03-011 | 11CN-03-011-W12, 11CN-03-011-W16 | 84, 112 | 0 |  |
| 11CN-03-018 | 11CN-03-018_b_w4 | 28 | 0 |  |
| 11CN-03-034 | 11CN-03-034-W14 | 98 | 0 |  |
| 11CN-03-040 | 11CN-03-040D8, 11CN-03-040-W8,<br>11CN-03-040-W12, 11CN-03-040-W16 | 8, 56, 84, 112 | 0 |  |
| 11CN-03-057 | 11CN-03-057D4 | 4 | 0 |  |
| 11CN-03-067 | 11CN-03-067D4 | 4 | 1 | EYB25_004707 |
| 11CN-03-067 | 11CN-03-067_b_d15 | 15 | 0 |  |
| 11CN-03-071_b_d0 | 11CN-03-071-D8 | 8 | 3 | EYB25_004793,<br>EYB25_007616 |
| 11CN-03-071_b_d0 | 11CN-03-071-D12 | 12 | 2 | EYB25_004798,<br>EYB25_004800 |
| 11CN-03-071_b_d0 | 11CN-03-071W18 | 126 | 7 |  |
| 11CN-03-072 | 11CN-03-072D4 | 4 | 2 |  |
| 11CN-03-072 | 11CN-03-072-D12 | 12 | 0 |  |
| 11CN-03-077 | 11CN-03-077_b_d11 | 11 | 0 |  |
| 11CN-03-087 | 11CN-03-087-D9 | 9 | 1 | EYB25_005954 |
| 11CN-03-103-D5 | 11CN-03-103-W4 | 23 | 0 |  |
| 11CN-03-122 | 11CN-03-122-W8 | 56 | 0 |  |
| 11CN-03-138 | 11CN-03-138-D11, 11CN-03-138_b_d35 | 11, 35 | 0 |  |
| 11CN-03-138-S | 11CN-03-138_s_m8 | 240 | 0 |  |
| 11CN-03-152 | 11CN-03-152_b_d14 | 14 | 2 | EYB25_000740,<br>EYB25_004486 |
| 11CN-03-159 | 11CN-03-159-D10 | 10 | 1 |  |
| 11CN-03-159 | 11CN-03-159-W12 | 84 | 1 |  |
| 11CN-03-162_s_d0 | 11CN-03-162-S-W24 | 168 | 3 | EYB25_002369,<br>EYB25_003095,<br>EYB25_003335 |
| **Blood** | **Skin** |  |  |  |
| 11CN-03-044 | 11CN-03-044-S | 0 | 0 |  |
| 11CN-03-046 | 11CN-03-046-S | 0 | 0 |  |
| 11CN-03-050 | 11CN-03-050-S | 0 | 0 |  |
| 11CN-03-057 | 11CN-03-057-S | 0 | 3 |  |
| 11CN-03-071 | 11CN-03-071-S | 0 | 2 |  |
| 11CN-03-075 | 11CN-03-075-S | 0 | 0 |  |
| 11CN-03-078 | 11CN-03-078-S | 0 | 0 |  |
| 11CN-03-085 | 11CN-03-085-S | 0 | 1 |  |
| 11CN-03-099 | 11CN-03-099_s_d0 | 0 | 0 |  |
| 11CN-03-102 | 11CN-03-102-S | 0 | 0 |  |
| 11CN-03-104 | 11CN-03-104_s_d0 | 0 | 0 |  |
| 11CN-03-108 | 11CN-03-108_s_d0 | 0 | 0 |  |
| 11CN-03-113 | 11CN-03-113_s_d0 | 0 | 1 |  |
| 11CN-03-115 | 11CN-03-115_s_d0 | 0 | 1 |  |
| 11CN-03-116 | 11CN-03-116_s_d0 | 0 | 1 |  |
| 11CN-03-122-W8 | 11CN-03-122-SW8 | 0 | 0 |  |
| 11CN-03-123 | 11CN-03-123_s_d0 | 0 | 0 |  |
| 11CN-03-133 | 11CN-03-133-S | 0 | 10 |  |
| 11CN-03-138 | 11CN-03-138-S | 0 | 0 |  |
| 11CN-03-140 | 11CN-03-140_s_d0 | 0 | 0 |  |
| 11CN-03-146 | 11CN-03-146-S | 0 | 0 |  |
| 11CN-03-148 | 11CN-03-148-S | 0 | 0 |  |
| 11CN-03-155 | 11CN-03-155_s_d0 | 0 | 1 |  |
| 11CN-03-160 | 11CN-03-160_s_d0 | 0 | 1 |  |
| 11CN-03-162 | 11CN-03-162_s_d0 | 0 | 0 |  |
| 11CN-03-166 | 11CN-03-166-S | 0 | 1 |  |
| 11CN-03-170 | 11CN-03-170-S | 0 | 1 |  |

We also observed that a subset of longitudinal isolates from 4 patients appear unrelated to the primary infecting isolate. In total, we observed 4 longitudinal blood isolates and 2 skin isolates (**Table 4**) that appeared unrelated to the initial blood isolate, or matched time point blood isolate, respectively. With SNP differences between these isolates and their initial or blood counterparts ranging from 1,072 to 2,810 (**Table 4**), it is likely that these cases represent novel strain introductions or pre-existing multi-strain infections, as opposed to a persistent infection of the same isolate over time.

**Table 4.** SNP differences in unrelated isolates taken from the same patient.

| Baseline | Longitudinal | Time (days) | SNPs | Related by phylogeny<sup>a</sup> |
| --- | --- | --- | --- | --- |
| 11CN-03-093 | 11CN-03-093-D8 | 8 | 2764 | FALSE |
| 11CN-03-093 | 11CN-03-093-D25 | 25 | 2810 | FALSE |
| 11CN-03-093 | 11CN-03-093_b_m6 | 180 | 2801 | FALSE |
| 11CN-03-132 | 11CN-03-132-D7 | 7 | 1427 | FALSE |
| 11CN-03-132 | 11CN-03-132-W12 | 84 | 1614 | FALSE |
| 11CN-03-138 | 11CN-03-138-M8 | 240 | 2721 | FALSE |
| **Blood** | **Skin** |  |  |  |
| 11CN-03-138-M8 | 11CN-03-138_s_m8 | 0 | 2142 | FALSE |
| 11CN-03-153 | 11CN-03-153-S | 0 | 1072 | FALSE |

<sup>a</sup>Related by phylogeny indicates whether these isolates co-localized in the phylogeny of isolates generated.

### Genomic variants associated with clinical outcomes

Given the range of clinical phenotypes and outcomes associated with isolates from both clades, we leveraged this variability to assess significant associations between clinical phenotypes and pathogen genetics. We took a genome wide association study (GWAS) approach to test for significance between loss-of-function (LoF) mutations and the clinical metadata suggestive of severe infections. Four patients with multiple infecting strains were excluded from this analysis. We saw LoF variants associate significantly (p<5×10^-6^, gemma score test) with poor clinical outcomes including mortality, relapse, IRIS, poor clinical response, and high baseline fungal burden (CFU/ml) (**Table 5**). These variants impacted genes with predicted roles regulating gene expression (methyltransferase, GAL4 transcription factor homolog), metabolism and fermentation (alcohol acetyltransferase, GTPase activating protein, malate synthase), cellular signaling (calcineurin-like phosphoesterase), protein processing (oligosaccharyltransferase, ubiquitin-protein ligase, exocyst complex component Sec3), and transport (HCO^3^- transporter, sugar transporter, major facilitator superfamily transporter). These isolates also varied in the MIC of itraconazole and amphotericin B, with a small number of isolates displaying notably elevated MIC values (**Figure S5**). While a GWAS did not identify any LoF variants significantly associated with higher MICs to these antifungals, there was very limited power in this analysis due to the low number of isolates with high MIC values. Genetic variation across a population may not only contribute to virulence but may also be selected for through environmental pressures. The variants identified here may give some advantage to survival within the environment of the human host and may be working in tandem with other forms of genetic variation, such as copy number, to elicit more severe clinical phenotypes.

**Table 5.** Variants significantly associated with clinical phenotypes.

| Phenotype<sup>a</sup> | Gene | P value | Gene function (Pfam) | Loss-of-function variant type |
| --- | --- | --- | --- | --- |
| Relapse | EYB25_006356 | 5.40E-07 | Metallophosphoesterase | Stop Gained |
| Relapse | EYB25_007086 | 1.39E-06 | Sec3 | Stop Gained |
| Relapse | EYB25_007995 | 1.54E-06 |  | Stop Gained |
| Relapse | EYB25_006573 | 2.15E-06 | Major facilitator superfamily | Frameshift |
| Mortality | EYB25_006471 | 4.44E-08 |  | Stop Gained |
| Mortality | EYB25_009504 | 2.50E-07 |  | Frameshift |
| IRIS | EYB25_004020 | 2.37E-12 | Methyltransferase | Frameshift |
| IRIS | EYB25_007265 | 8.05E-11 | AEX-3 | Stop Gained |
| IRIS | EYB25_009427 | 6.71E-10 | Alcohol acetyltransferase | Stop Gained |
| IRIS | EYB25_009447 | 4.68E-08 | Transcription factor | Frameshift |
| Poor clinical response | EYB25_006915 | 1.73E-18 |  | Stop Gained |
| Poor clinical response | EYB25_006356 | 2.93E-09 | Metallophosphoesterase | Stop Gained |
| Poor clinical response | EYB25_004962 | 3.84E-08 | Oligosaccharyltransferase | Stop Gained |
| Poor clinical response | EYB25_001778 | 4.96E-08 | Malate synthase | Stop Gained |
| Poor clinical response | EYB25_004898 | 4.22E-07 | HCO<sub>3</sub><sup>-</sup> transporter | Stop Gained |
| Poor clinical response | EYB25_008150 | 6.30E-07 | Sugar transporter | Frameshift |
| Poor clinical response | EYB25_006081 | 1.06E-06 | Haloacid dehalogenase hydrolase | Frameshift |
| Poor clinical response | EYB25_007654 | 2.32E-06 |  | Frameshift |
| High CFU | EYB25_007106 | 4.56E-06 | Ubiquitin transferase | Stop Gained |

<sup>a</sup>Significant variants ( $p < 5 \times 10^{-6}$ ) identified through multiple GWAS tests are combined in this table. Column 1 lists the phenotype that the variant is significantly associated with.

### Population diversity and regions under selection

To investigate larger scale genetic variation, we performed copy number variation analysis. We observed no evidence of chromosomal aneuploidy across these isolates but identified multiple genomic regions that appeared duplicated and deleted within this population. Many of these regional duplications and deletions appeared unique to individual isolates (**Figure 3**); however, common deletions were also detected. The most commonly deleted region encodes a predicted afadin- and alpha -actinin-binding region that is deleted across 141 isolates from the northern clade and 125 isolates from the southern clade. The largest number of isolates sharing a duplication across a single locus is 17, duplicated in isolates from the southern clade, this region encodes two reverse transcriptases, and a GAG-pre-integrase domain, likely corresponding to a retrotransposon integration. Of those deleted regions shared across more than 50 isolates, these encode genes predicted to function as transcription factors (6 genes), REDOX proteins (5 genes), protein kinases (4 genes), methyltransferases (3 genes), aspartic proteases (2 genes), dehydrogenases (2 genes), a cytochrome P450, a glycosyl hydrolase, a glycosyl transferase, a phospholipase, a phosphorylase, an adenylosuccinate lyase, an actin binding protein, a decarboxylase, an aminotransferase, an apoptosis regulating protein, an autophagy regulating protein, an arrestin, and a meiotic nuclear division protein. The range in functionality of genes lost across more than 50 isolates highlights the diversity within this clinical dataset. Of those duplicated regions shared across more than 2 isolates, these can be characterized by their predicted functions as heat shock and stress response genes, including heat shock protein 70, cell membrane and wall modulating genes, including a chitin synthase and a phosphoylcholine transferase, metabolic regulatory genes, including a cytochrome P450, a thiolase, a glycosyl hydrolase, an aspartyl protease, a citrate synthase, and gene expression regulatory genes, including a transcription factor and an mRNA capping protein. These results suggest that genes with roles in heat shock, cell wall structure, and metabolic response could provide an advantage to *T. marneffei* in this setting. We did not observe any duplications of the CYP51 ortholog within these isolates, suggesting alternate mechanisms may be responsible for the range in MIC values seen to itraconazole.

**Figure 3.**
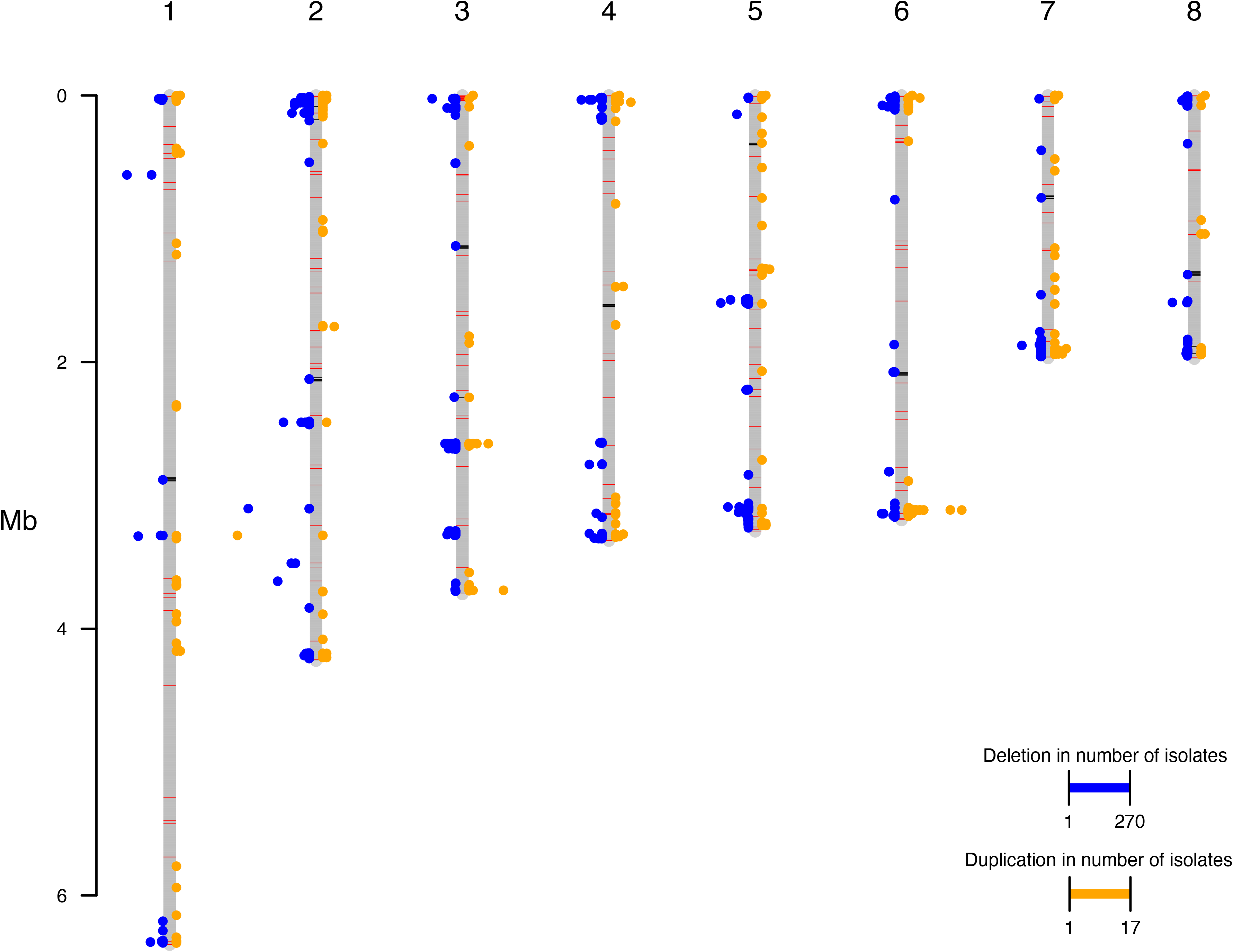
Frequencies of regional genomic duplications and deletions per chromosome. Frequency of deleted regions (blue points) and duplicated regions (orange points) assessed across all isolates, with frequency of deletion or duplication represented as distance from axis of the chromosomes (grey bars). 1000bp windows were used to assess copy number variation. GC content across chromosomes represented by black (GC 17-40%) and red (GC 50-57%) bars plotted on the grey chromosome bars.

To assess regions that may be under selection within these populations, we performed a population genetics analysis to identify rapidly evolving regions across these isolates. We noted strong signals present across many telomeric and sub-telomeric regions (**Figure 4a**), in tests for selective pressure (composite likelihood ratio, CLR) (48), nucleotide diversity (Pi), and population divergence (Dxy). To determine if these regions encode common functions, we performed enrichment analysis on all genes within the sub-telomeric regions. The sub-telomeric regions displaying strong signals for selective pressure were enriched (hypergeometric test, p<0.05) for genes predicted to play roles in thiamine diphosphate metabolism and energy derivation by oxidation of organic compounds (**Table S1**). When northern and southern clades were assessed independently via CLR analysis, non sub-telomeric regions exhibiting strong signals for selective pressure across both populations were enriched (hypergeometric test, p<0.05) for genes predicted to function in ion and organic anion transport, respiration, regulation of gene expression, and energy derivation by oxidation of organic compounds (**Table S1**). Non sub-telomeric regions displaying signals of population divergence between clades were enriched (hypergeometric test, p<0.05) for genes predicted to function in anaerobic respiration (**Table S1**).

**Figure 4.**
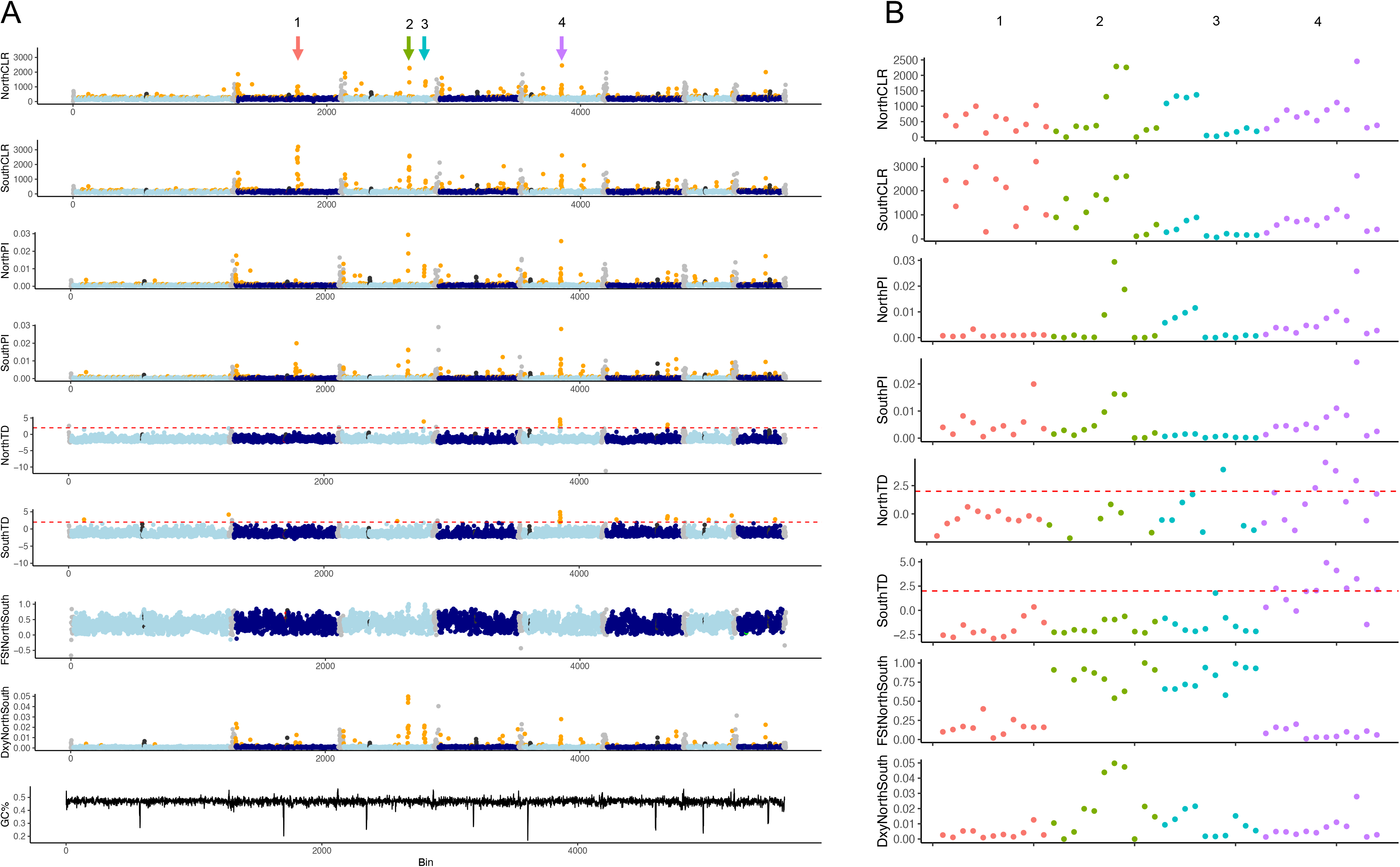
Genomic regions under selection across northern and southern clades. A) Signals of selection (CLR, TD), nucleotide diversity (Pi), and divergence (Fst, Dxy) across all chromosomes, per clade. B) Enlarged view of regions displaying elevated signals of selection, including region 1 (red), region 2 (green), region 3 (blue), and region 4 (purple).

To assess specific genes under positive selective pressures across these clades, we performed a composite likelihood ratio (CLR) analysis and found multiple regions with strong signals of selection across northern and southern clades, of which we will highlight four regions of interest (**Figure 4b**). Region 1 is under selection across isolates in both clades, but with relatively lower nucleotide diversity and CLR scores in the smaller northern clade (**Figure 4a**, region 1). This region shows lower levels of divergence between clades (Fst, Dxy), and appears to be rapidly evolving across all isolates with little clade specificity. This region encodes genes predicted to function as transcription factors (3 genes), protein kinases (3 genes), a ferric reductase, a methyltransferase, an adenylosuccinate lyase, and a phospholipase. Two additional regions displayed strong selective scores (CLR) across both clades in addition to elevated scores of divergence between clades (Fst, Dxy) (**Figure 4a**, regions 2 and 3). While region 2 shows high levels of nucleotide diversity for both clades, region 3 shows increased nucleotide diversity restricted to the northern clade, accompanied by an increased Tajima’s D score for this population. When region 2 and region 3 are interrogated further (**Figure 4b**), these two regions divide based on diversity within clade, with tracts of low diversity (Pi) within clades but high divergence (Dxy, Fst) observed between clades, indicating the potential for clade specific adaptations in these regions. Region 2 encodes genes predicted to function as a cytochrome P450 and a glycosyl hydrolase. Region 3 encodes genes predicted to function as a transcription factor, a protein kinase, an aspartyl protease, and a pyridoxal-dependent decarboxylase. To determine whether any of these signals may be due to a recent selective sweep, or balancing selection, we assessed scores for Tajima’s D. We identified one region that shared a strong signal in both our CLR analysis, and our Tajima’s D analysis (**Figure 4a**, region 4), indicating that this region may have recently undergone a selective sweep. We note that this region also shows an elevated Dxy score, but no increase in Fst between the two populations, indicating increased levels of divergence between the two populations at this region, accompanied by high levels of within-population diversity, as reflected in our measurements of nucleotide diversity (Pi) for each clade at this region. This region encodes 14 genes and multiple ankyrin repeats (EYB25_006920-006938) in total. The predicted functions of these genes include a transcription factor, an endoplasmic reticulum membrane protein, an arrestin, a meiotic nuclear division protein, a glycosyl transferase, a cell surface protein, an alcohol dehydrogenase, and an autophagy protein. These regions feature genes that may be playing important roles across and between clades, highlighting candidate genes for future functional analysis.

## Discussion

This large-scale genomic study of *T. marneffei* has begun to shed light on the population variability that exists within a large endemic area of talaromycosis. Phylogenetic analysis of 336 clinical isolates revealed a population of two major clades, consistent with a previous MLST study that identified geographically structured clades of *T. marneffei* (20). The clades representing both northern and southern Vietnam identified through MLST are recapitulated here using WGS based SNP phylogenies. Further, we calculate similar rates of LD decay for both clades individually as for together, indicating that recombination is taking place at similar levels within and between clades. Southern isolates show slightly increased clonality, harboring alleles with fewer SNPs and displaying significantly shorter terminal branches than isolates from the northern clade, highlighting the possibility of a more recent introduction.

Leveraging the clinical metadata available for these isolates, we investigated the possibility of clade-based differences with regards to clinical phenotypes and outcomes. We did not observe significant differences between the northern and southern clades with respect to 24-week mortality, rates of relapse, IRIS, clinical response to treatment, or MICs of itraconazole and amphotericin B. However, we detected significant associations between the southern clade and aspects of disease severity, including higher baseline fungal burden and presence of fever. In addition, we identified multiple loss-of-function variants significantly associated with mortality, relapse, IRIS, poor clinical response, and high baseline fungal burden. We noted that many of these clinical phenotypes may occur at low rates for the group sizes, likely impacting our ability to detect both significant differences between clades, and significant associations between variants and phenotypes. In addition, there are many confounding factors that contribute to these clinical outcomes, including host immune status, host response, timeliness of treatment, and antifungal treatment regimen. While we did not find that age, CD4 count, or ARV treatment status confounded associations between the southern clade and higher baseline fungal burden or presence of fever, larger studies of talaromycosis outcomes should be completed to provide increased power for estimations of clinical differences between clades to confirm these findings. Further work using larger group sizes for genotype-phenotype association studies will also be essential for building on our GWAS findings.

Applying a range of population genetics approaches with the variant data generated for these clinical isolates, we interrogated signals of selective pressure, increased nucleotide diversity, and population divergence for these clades. We observed shared signals of rapidly evolving regions present in northern and southern clades, with the strongest shared signals spanning regions containing genes with roles in gene expression regulation, iron sequestration, cellular signaling, and metabolism. Regions exhibiting signals of divergence between populations encoded genes with roles in metabolism and adaptation to environmental niches, sugar metabolism, transcriptional regulation, and protein cleavage. Of note, we observed strong signals of selection at the telomeric and sub-telomeric regions, in line with observations for other fungal pathogens such as *Cryptococcus neoformans* (15, 25). While sugar transporters are known to be enriched in these regions of *C. neoformans*, the subtelomeric regions displaying signals of selection in *T. marneffei* are enriched for genes involved in energy derivation by oxidation of organic compounds, and thiamine diphosphate metabolism.

The phylogenetic analysis of both longitudinal samples, and samples obtained from multiple body sites allowed for the inference of infections resulting from single, or multiple infecting isolates. We observed 6 instances in which longitudinal isolates obtained from a patient appeared unrelated to the primary infecting isolate, hinting at novel strain introductions, or the presence of multiple infecting strains. This is the first time this approach has been applied to *T. marneffei* to determine whether patients with protracted talaromycosis or infection recrudescence have been re-infected with novel isolates. This approach could also enable the identification of patients that have been chronically exposed, resulting in multiple contained primary infections that break up in the event of becoming immunocompromised. Sequencing of additional isolates at each time point would enable increased resolution to explore this possibility. The majority of patients that underwent longitudinal isolate collection showed continued infection with the basal isolate, in line with similar observations for *C. neoformans* (27, 49).

Combining whole genome sequencing data and patient data for this clinical cohort has enabled a better understanding of the relatedness of infecting isolates, within individual patients, and within the broader population. This approach enables identification of pathogen loci that appear under selection across both geographic clades, and highlights multiple genes that have been repeatedly, and independently, lost or duplicated across these isolates. Patient metadata allowed for association analyses that have given us insights into genetic signatures of the pathogen that are significantly associated with specific patient outcomes. Combining data types such as these, on an ever-larger scale, will afford a better understanding of the genetic basis of pathogenicity in fungal pathogens such as *T. marneffei*.

## Data Availability

All sequencing data are available via the accession PRJNA949141.

## Acknowledgments

The authors thank the Broad Institute Genomics Platform for sequencing isolates as part of this study.

## Funding

This project has been funded in part with Federal funds from the National Institute of Allergy and Infectious Diseases, National Institutes of Health, Department of Health and Human Services, under award U19AI110818 to the Broad Institute. This work was supported by the following NIH grants to TL (R01AI143409, R21AI162367, R21AI159397, U01AI169358, P30AI064518).

---

**Supplemental Table 1.** Functional enrichment of regions under selection across clades.

| Category | Test | GO Term | Adjusted p-value |
| --- | --- | --- | --- |
| Sub-telomeric | CLR | GO:0015980 | 0.04 |
| Sub-telomeric | CLR | GO:0042357 | 0.04 |
| Non sub-telomeric | CLR | GO:0015980 | 0.006 |
| Non sub-telomeric | CLR | GO:0045333 | 0.006 |
| Non sub-telomeric | CLR | GO:0015711 | 0.02 |
| Non sub-telomeric | CLR | GO:0006811 | 0.03 |
| Non sub-telomeric | CLR | GO:0010468 | 0.03 |
| Non sub-telomeric | Dxy | GO:0009061 | 0.05 |

**Supplemental Figure 1. Recombination rates per chromosome.**

Recombination rates (p/bp) as calculated by Ldhelmet, for a subset of 50 isolates.

**Supplemental Figure 2. Population structure and geographical clades.**

A) Population structure as determined by splitstree identifies two distinct clades, grouped by geography. B) Maximum likelihood phylogeny of patient isolates, with minimum inhibitory concentrations of itraconazole (ITR, blue) and amphotericin B (AMB, orange), visualized around the outer perimeter of the phylogeny.

**Supplemental Figure 3. Linkage disequilibrium decay.**

Linkage disequilibrium decay over 250 kb for northern (orange) and southern (blue) clades, as well as all isolates combined (grey).

**Supplemental Figure 4. Initial fungal burden and clearance rate by clade.**

A) Rate of clearance (Slope) by the infecting isolate clade. Displayed as −1(gradient). B) Blood Log_10_ CFU/mL (fungal burden) by infecting isolate clade.

**Supplemental Figure 5. Minimum inhibitory concentrations of itraconazole and amphotericin B.**

A) Histogram displaying the minimum inhibitory concentrations of itraconazole (ITR). B) Histogram displaying the minimum inhibitory concentrations of amphotericin B (AMB).

